# Endothelial tPA-dependent recruitment of microglia to vessels protects the blood-brain barrier after stroke in mice

**DOI:** 10.1101/2025.01.14.632943

**Authors:** Tamara Etuzé, Louis Fosset, Denis Vivien, Benoit D Roussel, Eloise Lemarchand

## Abstract

Thrombolysis with tissue-type plasminogen activator (tPA) remains the only pharmacological treatment for the acute phase of ischemic stroke. In this study, we hypothesize that endothelial tPA plays a key role in modulating the microglial response and maintaining blood brain barrier (BBB) integrity after stroke. Using a mouse model with conditional deletion for endothelial tPA (VeCad^Cre^ – tPA^Flox^) combined with a thrombotic stroke model and high-resolution imaging, we investigated the effects of endothelial tPA on vascular inflammation and microglia activation during the acute phase of stroke. Our results demonstrate that microglia-vessel contacts increase post-stroke. Notably, endothelial tPA deletion reduces vascular VCAM1 expression associated with decreased microglial activation and fewer microglia-vessel contacts. Following stroke, endothelial tPA deletion is associated with increased BBB permeability and heightened risk of haemorrhagic transformation. Collectively, these findings indicate that endothelial tPA mediates microglial recruitment to blood vessels, thereby exerting a protective effect on BBB integrity following ischemic stroke.

## Introduction

Ischemic stroke is one of the leading causes of death and disability worldwide. Currently, thrombolysis (using tissue type plasminogen activator; rtPA; or one of its mutant; tenecteplase) is the only pharmacological treatment available for the acute phase of ischemic stroke. This is often combined with endovascular thrombectomy, to limit brain damages caused by cerebral artery occlusion. Although rtPA is beneficial for its fibrinolytic action for the treatment of stroke, it has been shown that thrombolysis by rtPA can present adverse effects such as blood brain barrier (BBB) permeabilization and hemorrhagic transformations[1–3]. In addition, preclinical studies have shown that endogenous tPA, particularly neuronal tPA, also participates in the deleterious effects of tPA on the brain parenchyma [4,5]. Inflammation following stroke is well known to play a critical role in the progression of brain damage, with microglia which is the first immune cell type to respond to disruption in brain homeostasis and to produce important immune mediators. Accordingly, recent evidence indicates that the interaction between microglia and blood vessels plays a crucial role in maintaining BBB integrity during disease, as well as in neurovascular coupling[6,7]. For example, studies have shown that microglial P2RY12 is critical for interactions between microglia and BBB components[6,8]. However, the mechanisms of communication between microglia and blood vessels remain unclear. In this study, we postulate that endothelial tPA mediates the interaction between microglia and blood vessels during stroke, with an impact of the homeostasis of the BBB and inflammatory processes.

## Methods

### Animals

All animal procedures adhered to the European Council Directives 2010/63/EU and were approved by the local ethical committee of Normandy (CENOMEXA). Experiments were performed according to the Animal Research: Reporting of In Vivo Experiments (ARRIVE) guidelines and were conducted in a blinded manner from the experiments through to the analyses. All mice were housed in a room with automatically controlled temperature (21– 25 °C), relative humidity (45–65%), and light-dark (12–12h) cycles.

Four C57BL/6 mice (Janvier Labs) and 39 VeCadCre: tPAlox mice on a C57BL/6 background (Centre Universitaire de Ressources Biologiques, Caen, France), aged 12 to 16 weeks and weighting 25 to 30g, were used. Endothelial specific tPA knockout mice (VE-Cre^ΔtPA^) were generated by crossing mice with exon 3 of the Plat gene flanked by loxP sites with VE-Cadherin-Cre mice (B6.FVB-Tg(Cdh5-cre)7Mlia/J; # 006137; kind gift from F. Millat, Institute of Radioprotection and Nuclear Safety in Fontenay-aux-Roses, France), to obtain VE-Cre^ΔtPA^mice and VE-Cre^WT^mice.

### Thrombotic stroke model

Animals were anesthetized with 5% isoflurane in a gas mixture of 70% nitrous oxide and 30% oxygen. Temperature was maintained at 37+/-0.5°C using a warming blanket placed under the animal and a rectal probe during the procedure. The mice were placed in a stereotaxic device and maintained under anesthesia (1,5-2% isoflurane; 70% N_2_O/30% O_2_). A skin incision was performed between the right eye and ear, and the temporal muscle was dissociated. Then, a small craniotomy was performed on the parietal bone to expose the M1-M2 segments of the right middle cerebral artery (MCA). A Whatman filter paper strip, soaked in freshly prepared AlCl3 solution (40%, Sigma-Aldrich), was placed on the intact dura mater to cover the bifurcation of MCA and left in contact for 5 minutes. Cortical cerebral blood flow (CBF) in the MCA territory was measured using a laser Doppler flow probe (Oxford Optronix) positioned on the skull[9]. Sixteen mice were excluded due to inadequate arterial occlusion and absence of ischemic lesion.

### Magnetic Resonance Imaging (MRI)

MRI experiments were conducted on a Pharmascan 7T (Bruker, Germany). Mice were deeply anesthetized with 5% isoflurane and maintained with 1.5-2% isoflurane in 30% O2 / 70% N2O during the MRI acquisitions. High-resolution T2-weighted images were acquired to assess ischemic brain damage using a multislice multi-echo sequence with the following parameters: TE/TR 48.6 ms/3000 ms; slice thickness 0,5 mm; image size 256; average 2; field of view 17,92; scan time 3 min 12 sec. T1-weighted images were acquired to evaluate gadolinium extravasation, with the following parameters: TE/TR 15 ms/3 ms; slice thickness 9 mm; image size 256; average 3; field of view 17,92; scan time 4 min 1 sec 920 ms. T1-weighted acquisitions were performed both before and 5 minutes after gadolinium injection (Clariscan™Gé, GE Healthcare SAS). Gradient echo-planar imaging T2^*^-weighted sequences were used to detect hemorrhagic transformation, with the following parameters: TE/TR 8.706 ms/500 ms; Slice Thickness 0,5 mm; Image size 256; average 2; Scan time 3 min 12 sec. Lesion volumes, gadolinium extravasation and hemorrhagic volumes were determined using ImageJ® software on coronal brain sections obtained with MRI. Hemorrhagic transformation (HT) was scored as follows: (0) no HT; (1) small HT; (2) HT.

### Tissue processing

Anesthetized mice were transcardially perfused with cold heparinized saline. Brains were removed and post-fixed for 24h at 4°C with 4% paraformaldehyde. Then, brains were cryoprotected with 20% sucrose in PBS for 48h at 4°C before freezing in Epredia™ Cryomatrix™ embedding resin (Thermo Scientific). Blood was collected directly from the left ventricle and sodium citrate 0.129M was added to the collected blood fraction at a ratio of 1:9. Blood was centrifuged twice at 1,500g for 15min at room temperature to eliminate erythrocytes and platelets.

### Immunohistochemistry and immunofluorescence

Cryostat-cut sections (50 µm) were obtained and stored in freezing solution (ethylene glycol 30%, glycerol 20%, PBS 50%) at −n20°C. Sections were washed three times in PBS for 15min and incubated overnight at room temperature with primary antibodies diluted in 0.25% Triton X-100. Primary antibodies used: rat anti-VCAM1 (1:200, 553330 BD Pharmingen), goat anti-PODXL (1:200, AF1556 R&Dsystems), rabbit anti-Iba1 (1:500, ab178847 Abcam), rat anti-CD68 (1:1000; ab53444 Abcam). Primary antibodies were revealed by using Fab’2 fragments of Donkey anti-species linked to Alexa 488, Cy3 or Cy5 (1:500, Jackson ImmunoResearch). Secondary antibodies were incubated for 1h30min at room temperature.

### Quantification of microglia morphology and microglia-vessel interaction

Confocal images from coronal brain sections of defined areas (4 to 6 z-stack) were captured using Leica SP8 laser scanning confocal. Three-dimensional modeling was done on z-stacked images using Imaris7. Microglia morphology was quantified using 3DMorph[10], a MATLAB-based script that semi-automatically processes individual microglial morphology from three-dimensional (3D) data.

For the quantification of VCAM1 positive vessels and CD68+ microglia, binary masks were created for each channel using ImageJ and the function Image Calculator “AND” was used to measure the percentage of PODXL area covered by VCAM1 or the percentage of microglia covered by CD68. Results are expressed as % of VCAM1 positive vessels or % of CD68 positive signal in microglia.

For the quantification of microglia and vessel interaction, binary masks were created for PODXL+ and Iba1+ signals using ImageJ. The function Image Calculator “AND” was then used to measure the percentage of vessels (PODXL+) covered by microglia (Iba1+). Results are expressed as % of Iba1 positive signal (microglia) in contact with vessels. To investigate microglia recruitment to blood vessels, we measured the presence of microglia in a defined peri-vascular area corresponding to 2.9 µm around blood vessels (PODXL+). The function Image Calculator “AND” was then used to measure the percentage of the peri-vascular area that was positive for Iba1. Results are expressed as % of Iba1 positive signal in the peri-vascular area. Similar analyses were performed for the quantification of microglia in contact with VCAM1 positive and VCAM1 negative vessels. Individual datapoints represent the average of each z-stack measurement.

### Isolation of brain microvessels

Microvessels were isolated as previously described Boulay et al., 2015[11]. Brain tissue was placed in ice-cold HBSS containing 1M HEPES. Brain cortex was cut into 2-3mm pieces with scalpel, and tissue was dissociated with an automated Dounce homogenizer, performing 20 strokes at 400 rpm. Homogenate was centrifuged at 2,000g for 10 min at 4°C. The vessel pellet was manually and vigorously shaken for 1 min in ice-cold HBSS-HEPES containing 18% dextran, followed by centrifugation at 4,000g for 15 min at 4°C. The myelin in the upper layer and supernatant were discarded, and the vessels pellet was suspended in ice-cold HBSS-HEPES containing 1% BSA. The vessels solution was filtered (20µm pore size, hydrophilic nylon membrane, 47mm diameter, Millipore, NY2004700). The vessels were rinsed with HBSS-HEPES-BSA. Filter was recovered and gently shaken in HBSS-HEPES-BSA to detach the vessels. The vessels solution was centrifuged at 2,000g for 10 min at 4°C. Then, the vessels pellet was resuspended in HBSS-HEPES-BSA and transferred in a 1.5mL Eppendorf tube and centrifuged at 8000g for 10 min at 4°C. The vessels pellet was resuspended in 50µL RIPA lysis buffer (Thermo Scientific, 89900) with low binding tip, and vessels were dissociated with a 25G syringe. The mixture was centrifuged at 12,000g for 15 min at 4°C, and the supernatant was collected. Protein concentration was determined using the BCA protein assay (Thermo Scientific, 23225).

### Plasminogen-Casein Zymography

tPA enzymatic activity was assessed on 7,5% SDS-polyacrylamide gel containing 10 mg/mL bovine casein (ICN Biomedicals Inc., diluted in Tris 1.5 M pH 8.8) and 1 mg/mL of human plasminogen (Lys-Type, Calbiochem-Millipore, 528185). Samples (8 µg/well) were electrophoresed at 110V and 4°C for 3 hours 45 min. Gels were washed for 1 hour in 2.5% Triton X-100 and 30 minutes in H2O. Gels were incubated with 7.5 mg/mL glycine and 2.9 mg/mL EDTA for 2 hours at 37°C. Caseinolytic bands were visualized with Coomassie Blue staining.

### Statistical analysis

All statistical analysis and data representation were performed using GraphPad Prism 10.0 software and data are represented as mean ± SEM. Data were analyzed using unpaired t-test, one-way ANOVA (followed by Sidak’s multiple comparisons post hoc test) or Mann-Whitney for non-normally distributed variables. Kruskal Wallis test (followed by Dunnett’s multiple comparisons post hoc test) was used for non-continuous variables. Differences were considered statistically significant when *p* < 0.05.

## Results

### Microglia recruitment to blood vessels is mediated by endothelial tPA during the acute phase of ischemic stroke

First, vascular inflammation and microglial responses were assessed in the peri-infarct and contralateral areas 24 hours after the induction of a thrombotic stroke in mice (Figure 1A). We first revealed a significant increase of VCAM-1 expression in the peri-infarct area compared to the contralateral area (15.67-fold increase compared to contralateral, p=0.003; Figure 1B). In addition, expression of the lysosomal marker CD68 was significantly increased in microglia (1.85-fold increase compared to contralateral, p=0.0003; Figure 1C) and microglial ramifications were significantly decreased in the peri-infarct area compared to the contralateral area (1.94-fold decrease compared to contralateral, p<0.0001; Figure 1C). Vascular and microglial activation observed in our results are consistent with previous reports using different stroke models [12,13]. To investigate the communication between inflamed vessels and microglia, we measured microglial contacts with vessels. We observed a significant increase of microglial contacts with blood vessels in the peri-infarct area compared to the contralateral cortex (2.49-fold increase compared to contralateral, p=0.0061; Figure 1D-E). In addition, a significant increase of microglia in the perivascular area was observed (2.93-fold increase compared to contralateral, p=0.0008; Figure 1E) in both VCAM1-positive (7.57-fold increase compared to contralateral, p=0.0027, Figure 1F) and VCAM1-negative vessels (2.78-fold increase compared to contralateral, p=0.0015; Figure 1F). As tPA has been described to be released by endothelial cells following stroke, we then investigated the implication of endothelial tPA on microglia recruitment to blood vessels after stroke using a mouse model presenting a conditional deletion of endothelial tPA (Figure 2A). We first isolated microvessels from ipsilateral (red dots) and contralateral cortex (green dots) 24 hours after stroke in VE-Cre^WT^ and VE-Cre^ΔtPA^ mice (Figure 2B). Our data showed a significative decrease of endothelial tPA in the ipsilateral cortex compared to contralateral in VE-Cre^WT^ mice (7.70-fold decrease compared to contralateral, p <0.0001) and the lack of tPA in the VE-Cre^ΔtPA^ mice as expected (8.97-fold decrease compared to VE-Cre^WT^ mice, p<0.0001) evidencing an important release of endothelial tPA following stroke. Then, we investigated the impact of endothelial tPA deletion on vascular inflammation and microglia activation. We observed, 24 hours after stroke, a significant decrease in VCAM1 expression in peri-infarct vessels of VE-Cre^ΔtPA^ compared to VE-Cre^WT^ mice (2.21-fold decrease compared to in VE-Cre^WT^ group, p=0.0019; Figure 2C). In VE-Cre^ΔtPA^ mice, microglia also showed a significant decrease in CD68 expression (2.73-fold decrease compared to in VE-Cre^WT^ group, p=0.0014; Figure 2D) and an increase in microglial ramification (1.13-fold increase compared to in VE-Cre^WT^ group, p=0.0282; Figure 2D), indicating a reduced microglial reactivity in the peri-infarct area in VE-Cre^ΔtPA^ mice. In addition, we observed a decrease of microglia contacts with vessels (1.5-fold decrease compared to in VE-Cre^WT^ group, p=0.0187; Figure 2E-F*i*) and a decrease of the presence of microglia in the perivascular area (1.47-fold decrease compared to in VE-Cre^WT^ group, p=0.005; Figure 2F*i*) in both VCAM1-positive (2.73-fold decrease compared to in VE-Cre^WT^ group, p=0.0014; Figure 1F*ii*) and VCAM1-negative vessels (1.14-fold decrease compared to in VE-Cre^WT^ group, p=0.0282; Figure 1F*ii*). We also investigated if tPA was involved in microglial activation and recruitment to blood vessel in a model of neuroinflammation induced by LPS injection into the brain. We observed neither modification of VCAM1 expression, nor microglia activation and increase of microglial contacts with vessels in VE-Cre^ΔtPA^ compared to VE-Cre^WT^ mice in this model of neuroinflammation (Supplemental Figure 1). Altogether, these results suggest that endothelial cells can release tPA as a signal for the recruitment of microglia to blood vessels, specifically after thrombotic ischemic stress.

**Figure 1:**
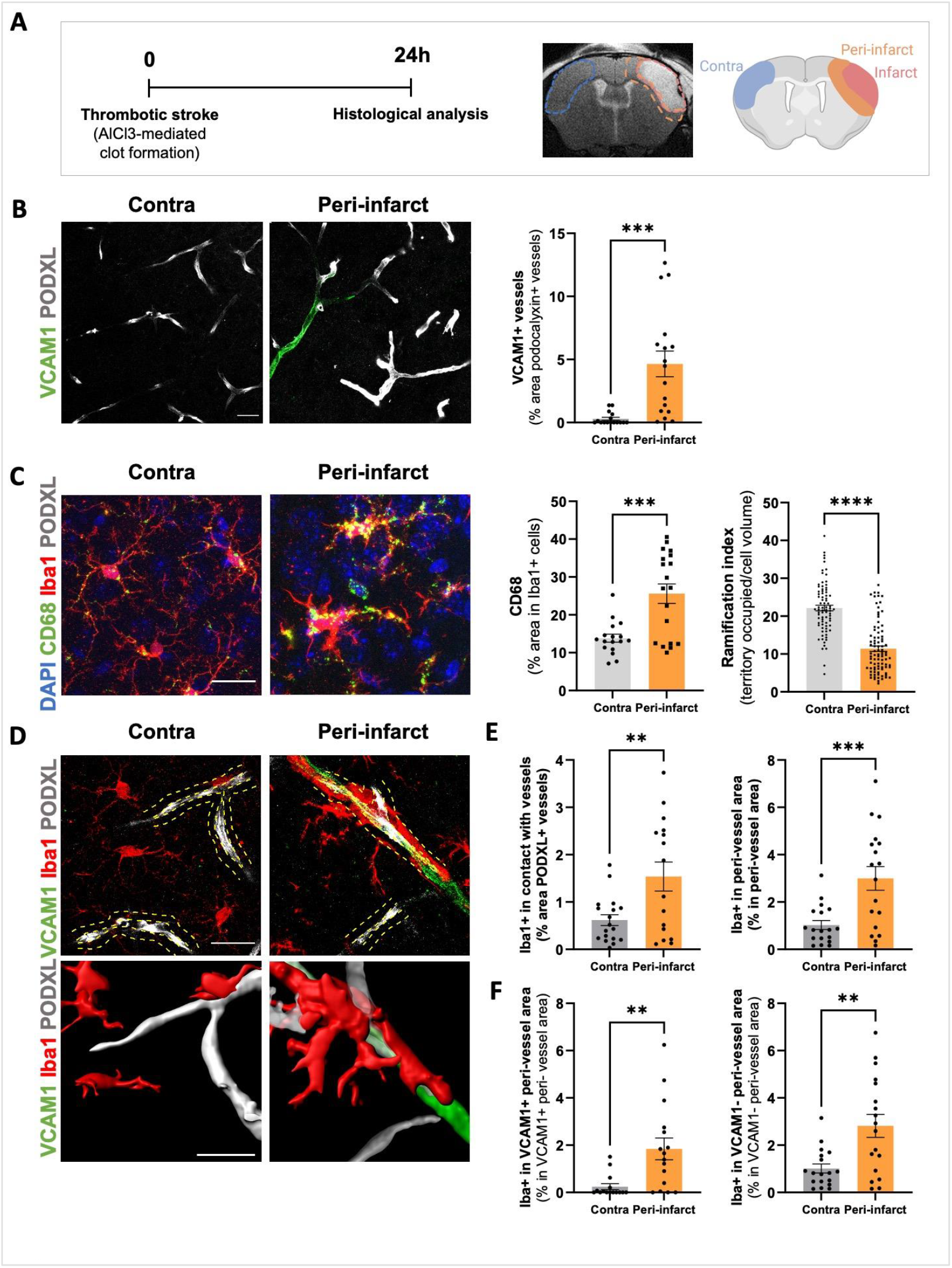
Vascular inflammation and microglial activation are associated with increased microglia-vessels interactions in the peri-infarct area after stroke. (A)Schematic representation of the experimental design. Created with BioRender.com. (B) Representative confocal images and quantification of VCAM1 positive vessels (podocalyxin, PODXL+) in contralateral and peri-infarct areas. Scale bar: 20 µm. ***p<0.001, two-tailed Student’s t-test, (N=4, n=16 contralateral and N=4, n=17 peri-infarct). (C) Representative confocal images of vessels (PODXL), microglia (Iba1) and CD68 staining in the contra and peri-infarct areas, quantification of CD68 expression in Iba1 positive cells (N=4, n=17 contralateral, N=4, n=19 peri-infarct) and quantification of microglia ramification index (territory occupied/cell volume), (N=4, n=80 contralateral and N=4, n=89 peri-infarct). DAPI labels cell nuclei. Scale bar: 20 µm. ^***^p<0.001, ^****^p<0.0001, two-tailed Student’s t-test, N=4. (D) Representative confocal images of vessels (PODXL), microglia (Iba1) and VCAM1 staining in the contra and peri-infarct areas. (E) Quantification of contact of microglia with vessels and quantification of microglia present in peri-vessels areas (dotted yellow lines represent peri-vessels areas). (F) Quantification of microglia in VCAM1 positive and VCAM1 negative peri-vessels areas. Scale bar: 20 µm. ^**^p<0.01, ^***^p<0.001 two-tailed Student’s t-test, (N=4, n=18 per group). Data are shown in mean ±SEM.

**Figure 2:**
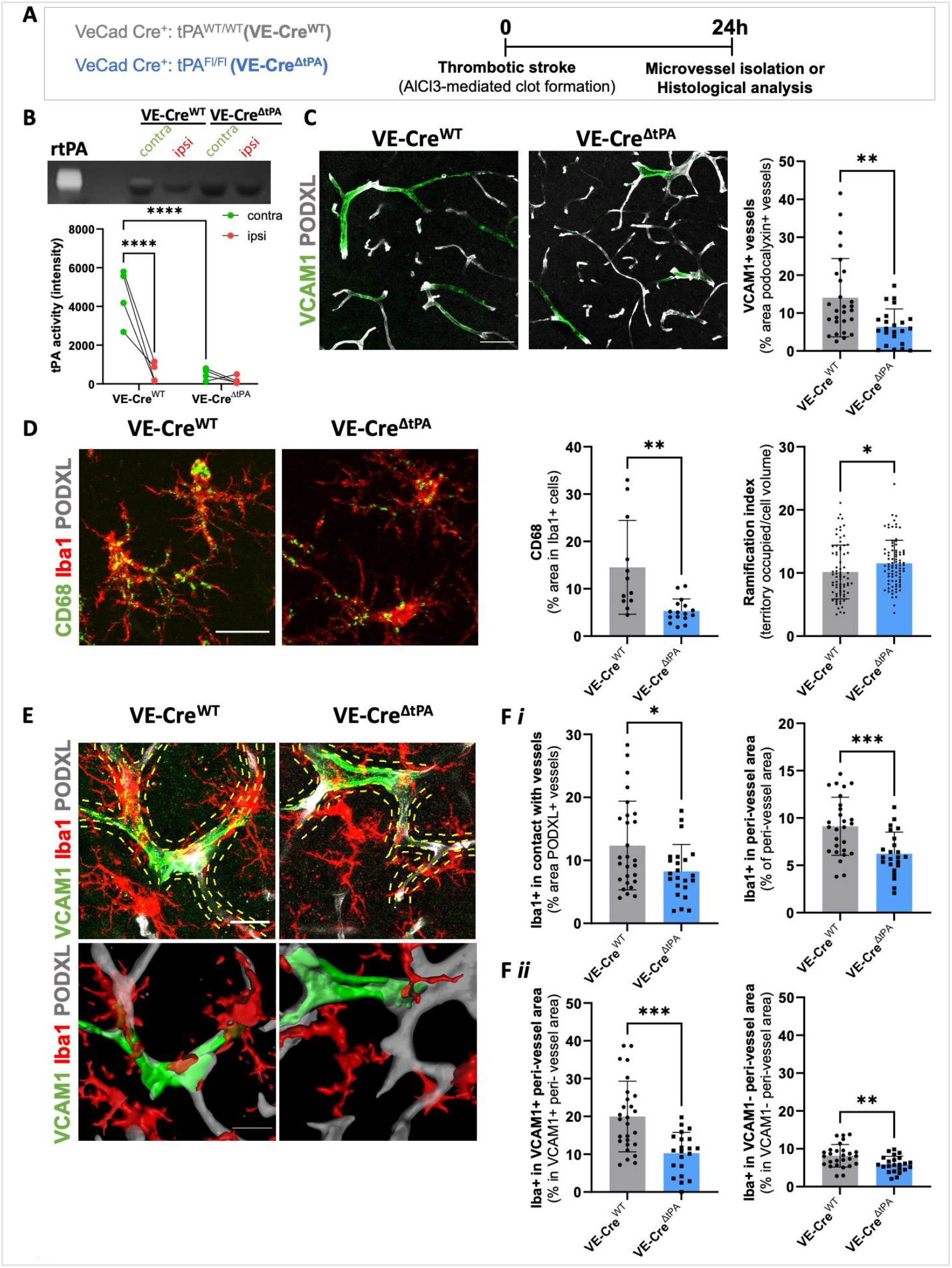
Endothelial tPA contributes to microglia recruitment to blood vessels after stroke. (A) Schematic representation of the experimental design. (B) tPA activity in isolated microvessels from ipsilateral (ipsi) and contralateral (contra) cortex 24h after stroke. ^****^p<0.0001, one-way ANOVA (N = 4 per group). (C) Representative confocal images and quantification of VCAM1 positive vessels (podocalyxin, PODXL+) in the peri-infarct area. Scale bar: 50 µm. ^**^p<0.01, two-tailed Student’s t-test (N=5, n=27 VE-Cre^WT^ and N=4, n=23 VE-Cre^ΔtPA^). (D) Representative confocal images of vessels (PODXL), microglia (Iba1) and CD68 staining in the peri-infarct area, quantification of CD68 expression in Iba1 positive cells (N=3, n=12 VE-Cre^WT^ and N=4, n=16 VE-Cre^ΔtPA^) and quantification of microglia ramification index (territory occupied/cell volume), (N=4, n=74 VE-Cre^WT^ and N=3, n=86 VE-Cre^ΔtPA^). Scale bar: 30 µm. *p<0.05, **p<0.01, two-tailed Student’s t-test. (E) Representative confocal images of vessels (PODXL), microglia (Iba1) and VCAM1 staining in the peri-infarct area, (F *i*) quantification of contact of microglia with vessels and quantification of microglia present in peri-vessels areas. (F *ii*) Quantification of microglia in VCAM1 positive and VCAM1 negative peri-vessels areas. Scale bar: 20 µm. ^*^p<0.05, ^**^p<0.01 ^***^p<0.001, two-tailed Student’s t-test (N=5, n=27 VE-Cre^WT^ and N=4, n=23 VE-Cre^ΔtPA^). Data are shown in mean ±SEM.

### Impact of endothelial tPA on BBB integrity and hemorrhagic transformation following ischemic stroke

As microglia interaction with blood vessels has been described to be critical in the maintenance of the BBB[7,8], we investigated the consequence of endothelial tPA deletion and altered microglia - blood vessels interaction on stroke outcome. We conducted a longitudinal study to monitor the evolution of brain lesion, gadolinium extravasation into the brain and hemorrhagic transformation between 24h and 5 days post-stroke using MRI (Figure 3A-B). Brain lesion volumes did not differ in the presence or absence of endothelial tPA (14.7mm^3^ *versus* 12.9 mm^3^ in VE-Cre^ΔtPA^ group at 24h, p=0.6670 and 7.3 mm^3^ *versus* 6.5 mm^3^ in VE-Cre^ΔtPA^ group at 5 days, p=0.9129; Figure 3C). However, gadolinium extravasation into the brain was significantly increased 5 days after stroke in VE-Cre^ΔtPA^ mice compared to VE-Cre^WT^ mice (3.75mm^3^ *versus* 5.70mm^3^ in VE-Cre^ΔtPA^ group at 5 days, p=0.0330; Figure 3D). This was accompanied by a surprisingly significant increase of hemorrhagic transformation scores and volumes in VE-Cre^ΔtPA^ mice (0.03mm^3^ *versus* 0.13mm^3^ in VE-Cre^ΔtPA^ group at 5 days, p=0.0080; Figures 3E-F). Altogether, these results tend to demonstrate that endothelial tPA plays a crucial role in maintaining BBB integrity and limiting hemorrhagic transformation after stroke.

**Figure 3:**
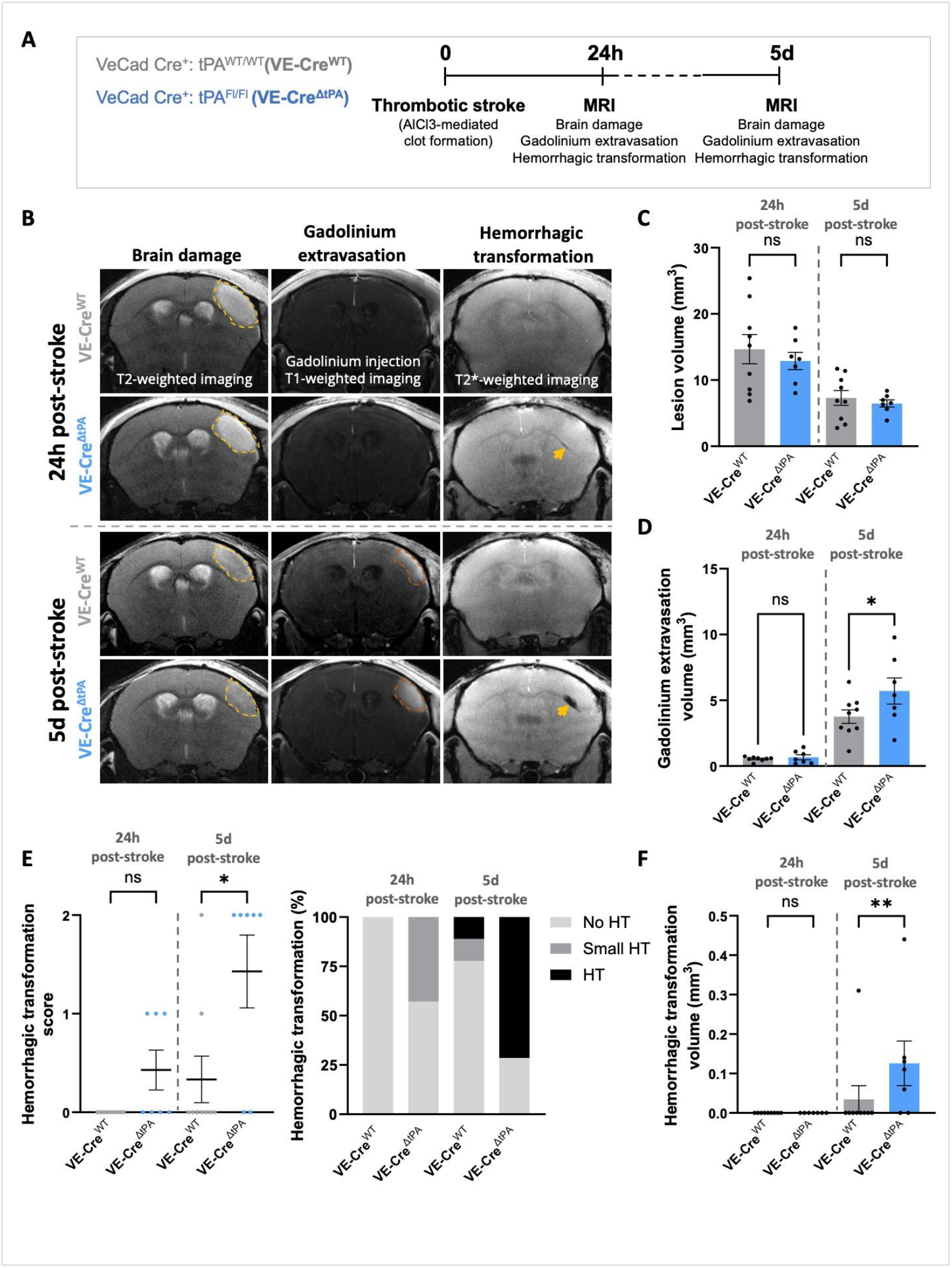
Endothelial tPA limits BBB permeabilization and hemorrhagic transformation 5 days after stroke. (A) Schematic representation of the experimental design. (B) Representative T2-weighted, T1-weighted and T2*-weighted images at 24h and 5 days after stroke. Edema are delineated by a discontinuous yellow line and hemorrhagic transformation (HT) is shown by yellow arrow. Quantification of (C) lesion volumes and (D) gadolinium extravasation volumes. *p<0.05, Ordinary one-way ANOVA. (E) HT score (no hemorrhagic transformation (0), small hemorrhagic transformation (1), hemorrhagic transformation (2)), distribution of HT score per group and (F) HT volume. ^*^p<0.05, ^**^p<0.01, Kruskal-Wallis test. Data are shown in mean ±SEM (N=9 VE-Cre^WT^ and N=7 VE-Cre^ΔtPA^).

## Discussion

Although thrombolysis with rtPA is beneficial for its fibrinolytic action for the treatment of stroke, it has been shown to have adverse effects such as BBB permeabilization and hemorrhagic transformations. These effects are in part due to an altered inflammatory response induced by rtPA[1,3]. However, the role of endogenous tPA in modulating the inflammatory response, particularly its influence on microglial activation, remains less understood. Our aim was to address this issue by investigating the impact of endothelial tPA on microglial responses and its subsequent effects on BBB permeabilization and hemorrhagic transformations.

We demonstrated that endothelial tPA plays a crucial role in recruiting microglia to blood vessel in the peri-infarct area during the acute phase of ischemic stroke. However, endothelial tPA does not appear to be involved in microglia – vessel communication in a model of neuroinflammation, demonstrating a strong specificity for ischemic stroke. Previous studies have shown that microglial P2RY12 is essential for microglial migration and interaction with blood vessels[6,7]. We previously reported that endothelial tPA is released immediately after stroke onset to mediate endogenous fibrinolysis and contribute to neurovascular coupling[14,15]. Here, we further demonstrate that endothelial cells release tPA to induce microglial recruitment to blood vessels following ischemic stress.

In addition, our results indicate that microglial recruitment to blood vessels after stroke helps to limit BBB permeability. Clinical and preclinical studies have demonstrated the pro-hemorrhagic effects of thrombolysis with exogenous rtPA[2,3,16]. In this study, we show a protective effect of endogenous tPA against hemorrhagic transformation and BBB permeabilization following stroke. These results may be attributed to the significantly higher doses of recombinant tPA injected for thrombolysis compared to the localized release of endothelial tPA in response to ischemic stress. In conclusion, we demonstrate that endothelial tPA serves as a key modulator of the interaction between microglia and endothelial cells following stroke.

## Acknowledgements

This work was supported by grants from the Ministère de l’Enseignement Supérieur et de la Recherche and INSERM (French National Institute for Health and Medical Research; HCERES U1237-2023/2028), tPAMicroglia (ANR-21-CE14-0039 [E.L.]) and by the Fondation pour la Recherche Médicale (ARF202005011926 [E.L.]).

## Authorship

### Contribution

T.E, L.F., E.L. performed the experiments. E.L., T.E., generated the figures and analyzed the data E.L., D.V., B.D.R. designed the study. E.L., D.V., B.D.R., T.E. interpreted the data and wrote the manuscript.

## References

1. Wang R, Zhu Y, Liu Z, Chang L, Bai X, Kang L, et al. Neutrophil extracellular traps promote tPA-induced brain hemorrhage via cGAS in mice with stroke. Blood. 2021;138:91–103.

2. Goncalves A, Su EJ, Muthusamy A, Zeitelhofer M, Torrente D, Nilsson I, et al. Thrombolytic tPA-induced hemorrhagic transformation of ischemic stroke is mediated by PKCβ phosphorylation of occludin. Blood. 2022;140:388–400.

3. Shi K, Zou M, Jia DM, Shi S, Yang X, Liu Q, et al. TPA Mobilizes Immune Cells That Exacerbate Hemorrhagic Transformation in Stroke. Circ Res. 2021;128:62–75.

4. Wang YF, Tsirka S, Strickland S, Stieg PE, G Ss, Lipton SA. Tissue plasminogen activator (tPA) increases neuronal damage after focal cerebral ischemia in wild-type and tPA-deficient mice. Nat Med. 1998;4:228–31.

5. Nicole O, Docagne F, Ali C, Margaill I, Carmeliet P, MacKenzie ET, et al. The proteolytic activity of tissue-plasminogen activator enhances NMDA receptor-mediated signaling. Nat Med. 2001;1:59–64.

6. Bisht K, Okojie KA, Sharma K, Lentferink DH, Sun YY, Chen HR, et al. Capillary-associated microglia regulate vascular structure and function through PANX1-P2RY12 coupling in mice. Nat Commun. 2021;12.

7. Császár E, Lénárt N, Cserép C, Környei Z, Fekete R, Pósfai B, et al. Microglia modulate blood flow, neurovascular coupling, and hypoperfusion via purinergic actions. J Exp Med. 2022;219.

8. Lou N, Takano T, Pei Y, Xavier AL, Goldman SA, Nedergaard M. Purinergic receptor P2RY12-dependent microglial closure of the injured blood-brain barrier. Proc Natl Acad Sci U S A. 2016;113:1074–9.

9. Le Behot A, Gauberti M, De Lizarrondo SM, Montagne A, Lemarchand E, Repesse Y, et al. GpIbα-VWF blockade restores vessel patency by dissolving platelet aggregates formed under very high shear rate in mice. Blood. 2014;123:3354–63.

10. York EM, Ledue JM, Bernier LP, Macvicar BA. 3Dmorph Automatic Analysis of Microglial Morphology in Three Dimensions From Ex Vivo and in Vivo Imaging. eNeuro. 2018;5:1–12.

11. Boulay AC, Saubaméa B, Declèves X, Cohen-Salmon M. Purification of mouse brain vessels. J Vis Exp. 2015;2015:1–8.

12. Gauberti M, Montagne A, Marcos-Contreras OA, Le Béhot A, Maubert E, Vivien D. Ultra-sensitive molecular MRI of vascular cell adhesion molecule-1 reveals a dynamic inflammatory penumbra after strokes. Stroke. 2013;44:1988–96.

13. Otxoa-de-Amezaga A, Miró-Mur F, Pedragosa J, Gallizioli M, Justicia C, Gaja-Capdevila N, et al. Microglial cell loss after ischemic stroke favors brain neutrophil accumulation. Acta Neuropathol. 2019;137:321–41.

14. Furon J, Yetim M, Pouettre E, Martinez de Lizarrondo S, Maubert E, Hommet Y, et al. Blood tissue Plasminogen Activator (tPA) of liver origin contributes to neurovascular coupling involving brain endothelial N-Methyl-D-Aspartate (NMDA) receptors. Fluids Barriers CNS. 2023;20:1–17.

15. Furon J, Lebrun F, Yétim M, Levard D, Marie P, Orset C, et al. Parabiosis Discriminates the Circulating, Endothelial, and Parenchymal Contributions of Endogenous Tissue-Type Plasminogen Activator to Stroke. Stroke. 2024;55:747–56.

16. Thiebaut AM, Gauberti M, Ali C, Martinez De Lizarrondo S, Vivien D, Yepes M, et al. The role of plasminogen activators in stroke treatment: fibrinolysis and beyond. Lancet Neurol. 2018;17:1121–32.

