## Supplemental Figure 1 for "Endothelial tPA-dependent recruitment of microglia to vessels protects the blood-brain barrier after stroke in mice"

**Supplemental Figure 1**

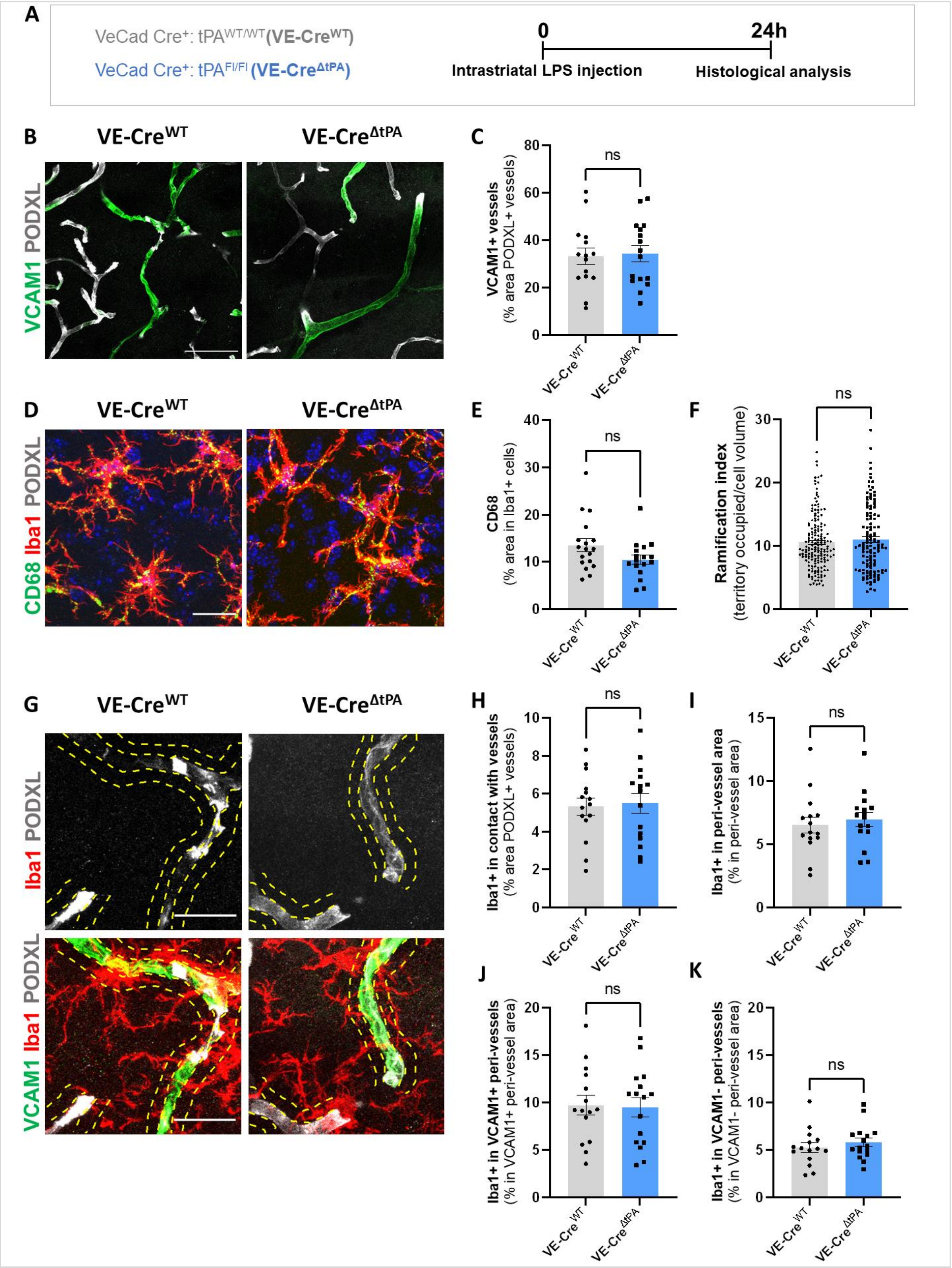

**Supplemental Figure 1: Endothelial tPA does not modify VCAM1 expression and microglial activation after LPS-induced neuroinflammation.**

(A) Schematic representation of experimental design. (B) Representative confocal images and (C) quantification of VCAM1 positive vessels (podocalyxin, PODXL+) in the striatum 24h after LPS (0.5 ug) injection. DAPI labels cell nuclei. Scale bar: 50 µm. Two-tailed Student's t-test, (N=3, n=15 VE-Cre$^{WT}$ and N=3, n=16 VE-Cre$^{\Delta tPA}$). (D) Representative confocal images of vessels (PODXL), microglia (Iba1) and CD68 staining in the striatum, (E) quantification of CD68 expression in Iba1 positive cells (N=3, n=15 VE-Cre$^{WT}$ and N=3, n=16 VE-Cre$^{\Delta tPA}$) and (F) measurement of microglia ramification index (territory occupied/cell volume), (N=3, n=169 VE-Cre$^{WT}$ and N=3, n=129 VE-Cre$^{\Delta tPA}$). (G) Representative confocal images of vessels (PODXL), microglia (Iba1) and VCAM1 staining in the striatum (dotted yellow lines represent peri-vessels areas) and quantification of (H) contact of microglia with vessels. Quantification of (I) microglia present in peri-vessels areas and (J) microglia in VCAM1 positive and (K) VCAM1 negative peri-vessels areas. DAPI labels cell nuclei. Scale bar: 20 µm. Two-tailed Student's t-test. Data are shown in mean ±SEM. (N=3, n=15 VE-Cre$^{WT}$ and N=3, n=16 VE-Cre$^{\Delta tPA}$).
